# genomeRxiv: a genome-based identification web server that bridges the gap between taxonomy and strain typing

**DOI:** 10.64898/2026.10.09.757156

**Authors:** Reza Mazloom, Rituraj Sharma, Janet S.H. Lorv, Sehgeet Kaur, Parul Sharma, C. Titus Brown, N. Tessa Pierce-Ward, Leighton Pritchard, Lenwood S. Heath, Boris A. Vinatzer

## Abstract

Precise identification depends on highly resolved classification. Therefore, whole genome-based strain typing schemes have been implemented for identification of the most important microbial human pathogens. However, there are no dedicated typing schemes for the majority of prokaryotic human, animal, and plant pathogens, or for non-pathogenic microbes, often limiting their identification to taxonomic classification tools that have the species as the smallest unit. To address this deficiency, we created the *genomeRxiv* web server which contains over 350,000 genomes and leverages the average nucleotide identity (ANI)-based Life Identification Number^®^ (LIN^®^) approach, allowing users to identify prokaryotic genomes in real time from approximately genus-rank to near strain-level resolution. LINs are overlaid with circumscriptions of species and infraspecies classes, integrating taxonomy with strain typing. Metadata and distance trees are visualized to complement inferred phylogenetic relationships with epidemiological and ecological information. A command line tool is provided for users to upload *k*-mer sketches of unpublished genomes and associated metadata, facilitating epidemiological investigations without the need to share complete DNA sequences. In conclusion, *genomeRxiv* constitutes a much-needed resource for precise, rapid identification of the many prokaryotes not included in any pathogen-specific typing scheme.

## Introduction

Today’s connected world, characterized by worldwide travel and trade, dramatically increases the risk of human, animal, and plant epidemics and pandemics. Antibiotic-resistant bacterial pathogens spread on asymptomatic carriers across the world^1^, lineages of animal pathogens emerge and spread through legal or illegal animal trade^2^, and plant pathogens are exchanged between continents on asymptomatic plants leading to new emerging crop diseases^3^. To investigate such emerging pathogen strains, to unravel their dissemination patterns, and to reduce the risk of their further spread, we need to be able to precisely identify them. We also require precise identification of microbes presumed to be innocuous, including those intended for biotechnological applications or for release in agricultural fields to stimulate plant growth or control diseases, to assess the risk that they may cause harm to humans, animals, and plants^4^.

The resolution of identification depends on three main factors: the analytical method that is deployed, the reference database that is available, and the classification scheme that is applied. To investigate outbreaks by bacterial pathogens, whole genome sequencing has become the analytical method of choice as it provides the most comprehensive genetic information possible^5^. The number of genome sequences in databases is also growing exponentially^6,7^. However, when using hierarchical Linnean taxonomy, the resolution of identification is limited to the species rank, even when using whole genome-based tools such as the Genome Taxonomy Database tool kit (GTDB-Tk)^8^ or TYGS^9^. Therefore, whole genome-based strain typing methods that use the single genome (or classes with a breadth very close to a single genome) as the smallest unit are deployed for epidemiological investigations, allowing for much more precise identification. Examples include GenomeTrakr, which uses single nucleotide polymorphisms (SNPs)^10^, and Enterobase^11^ and BIGSdb-Pasteur^12^, which use core-genome Multilocus Sequence Typing (cgMLST)^13^. These are robust systems with sufficient resolution to track disease outbreaks. However, they do not provide a standardized, hierarchical taxonomy. Therefore HierCC^14^ and LIN codes^15^ were developed to assign cgMLST-based sequence types (STs) to hierarchical infraspecies classes based on the number of shared alleles, informing about phylogenetic relationships and facilitating communication about disease outbreaks. These systems provide high resolution but must be set up specifically for each pathogenic species (or species complex) and so are currently only available for a few of the most important pathogens. For the majority of animal and plant pathogens and non-pathogenic prokaryotes, there is no comparable system to provide strain-level identification. This hinders epidemiological investigations and the establishment of effective regulatory policies and procedures, which often require within-species classification and identification^16^.

To address these issues, we developed the Life Identification Number^®^ (LIN^®^) system^17,18^. LINs provide a highly resolved hierarchical system of taxonomy, which uses the individual genome sequence instead of the species as the basic unit of classification. In place of the small number of Linnean ranks, the LIN system uses a much higher number of ranks based exclusively on quantitative measures of genome similarity^19^. In its current implementation, we use average nucleotide identity (ANI) to measure genome similarity. ANI is widely used to circumscribe prokaryotic species at or near the 95% threshold^20,21^, but we and others have shown that ANI can be informative about phylogenetic relationships from approximately genus rank (∼75-80% ANI) to almost single strain level (∼99.95%)^18,22–25^. In the current LIN system (**Fig. 1**), the highest rank, labeled A, is assigned at the 70% ANI threshold and the lowest rank, labeled T, is assigned at 99.999% ANI threshold, spanning the genus to strain interval. The last three thresholds from R (99.975% ANI) to T (99.999% ANI) are also a result of differences in gene content and genome sequence quality. Differences between LINs at these last three positions are thus more difficult to interpret but they do have the merit of providing a unique identifier to each distinct genome sequence.

**Fig. 1.**
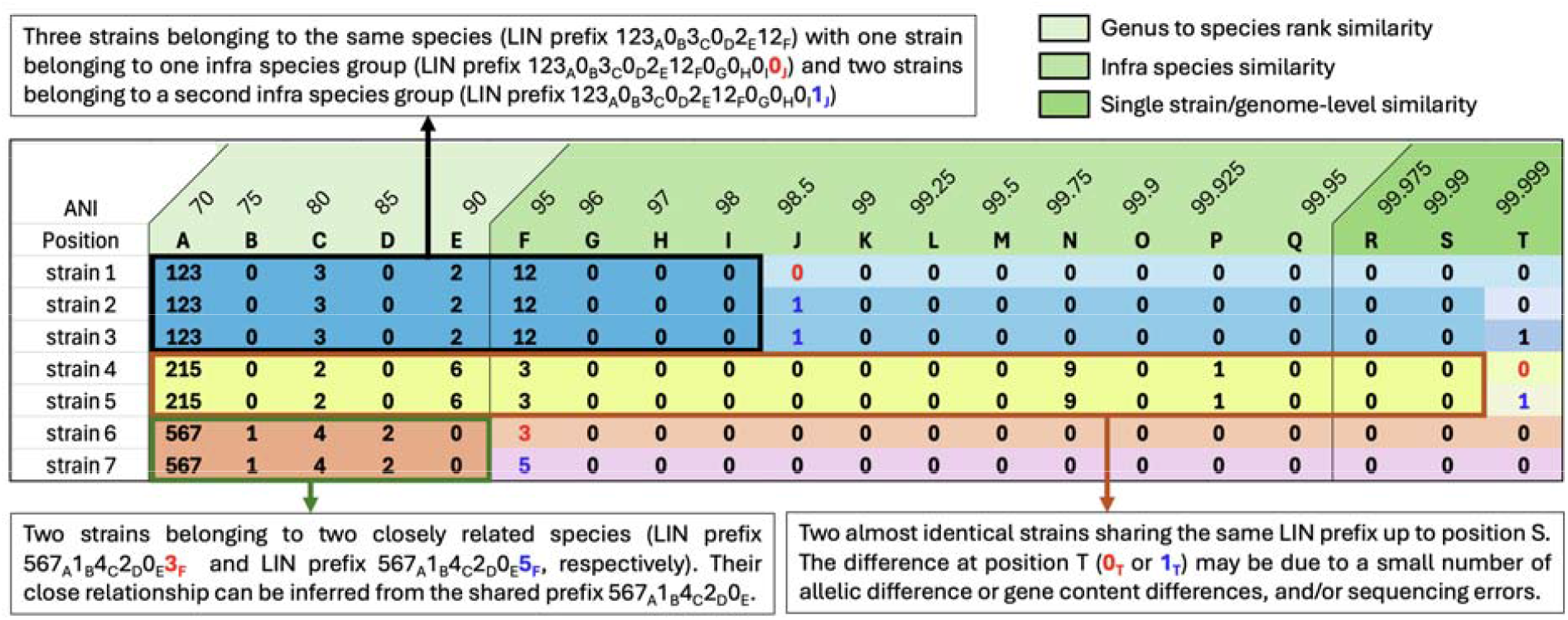
Schematic of LINs and LINgroups. LINs are assigned to each individual genome whereby the length of the LIN prefix shared by genomes indicates how similar genomes are to each other. LINs thus function as a hierarchical strain typing scheme. Text boxes in the figure provide examples of genomes sharing LIN prefixes of different length and how to interpret these shared prefixes. Any group of genomes sharing a LIN prefix of any length is called a LINgroup.

Alignment-based phylogenetic approaches are computationally demanding and relatively slow, but a LIN can be assigned to any prokaryotic genome in seconds to minutes^26^ by combining computationally efficient *k*-mer-based FracMinHash implemented in sourmash^27^ with slower but more precise alignment-based ANIb^28^. LINs thus embody a fast and highly resolved strain typing scheme applicable to all prokaryote genomes. The current LIN system has a high number of thresholds, meaning that almost any monophyletic group below genus rank, be it a species complex, species, or infraspecies class (e.g. phylotype, phylogroup, clade, genomovar, or clonal lineage) can in principle be precisely circumscribed using LIN positions (the LIN prefix) shared by its member genomes (**Fig. 1**). In other words, LINs bridge the gap between strain typing and taxonomy because LINs assigned to individual strains/genomes can be used as “coordinates” to circumscribe groups of related genomes, *i.e.*, taxa (similarly to how GPS coordinates can be used to circumscribe a geographic or political region). We refer to groups of strains that share a LIN prefix as a LINgroup. To the extent that a LINgroup consists of genomes of organisms with shared properties, identifying a query genome as a member of a LINgroup also predicts its likely phenotype, for example, high virulence or antibiotic resistance.

We previously established the *LINbase* web server to use LINs for precise prokaryotic identification^29^. However, because of the algorithm used for LIN assignment^26^, LIN assignment did not scale sufficiently to assign LINs to the growing number of public prokaryotic genomes. We constructed the *genomeRxiv* web server (genomerxiv.cs.vt.edu) to solve this performance issue, and to provide additional features facilitating data sharing for outbreak detection and scientific networking. These include enabling users to classify and share *k*-mer sketches in place of genome sequences (addressing restrictions on sequence sharing due to the Nagoya protocol^30,31^, intellectual property considerations, or other privacy concerns), and provision of visualizations that allow users to place sequenced genomes into evolutionary, epidemiological, and ecological contexts by integrating genome distance trees with available metadata. In summary, *genomeRxiv* leverages LINs to combine strain typing with taxonomic identification to precisely classify and identify prokaryotes, including the majority of species for which specific strain typing schemes are not available.

## Results

### Overview of the *genomeRxiv* web server

The *genomeRxiv* web server components and functions are summarized in **Fig. 2**. Briefly, the *genomeRxiv* web server was designed to classify any prokaryotic genome assembly in near-real time, either using the web interface or by connecting via the *BrookLIN* command line tool. Alternatively to providing genome sequences, users can also query with FracMinHash sketches, computed offline using sourmash^27^ or in real-time in the user’s browser, allowing for private queries in which the genome sequence never leaves the user’s computer. The primary result returned to the user is a temporary LIN that reflects the query’s similarity to all genomes in the *genomeRxiv* database with which it shares at least 70% ANI. In other words, the temporary LIN is the output of the strain typing function of *genomeRxiv.* The temporary LIN is then compared to stored LIN prefixes associated with circumscribed, named species and other named monophyletic groups to place the query sequence taxonomically, *i.e.*, the output of the taxonomic identification function of *genomeRxiv*. After the identification step (or, alternatively, when browsing or searching genomes and LINgroups in the database using keywords or LINs), users can visualize available source data, collection dates, geographic data, and distance trees for any genomes and signatures of LINgroups to which their query belongs in order to gain epidemiological, ecological, and evolutionary insights. Finally, the command line tool *BrookLIN* allows users to perform these *genomeRxiv* functions on batches of genomes or sourmash signatures, and also to share signatures and associated metadata by uploading them to *genomeRxiv,* making them discoverable by other users.

**Fig. 2.**
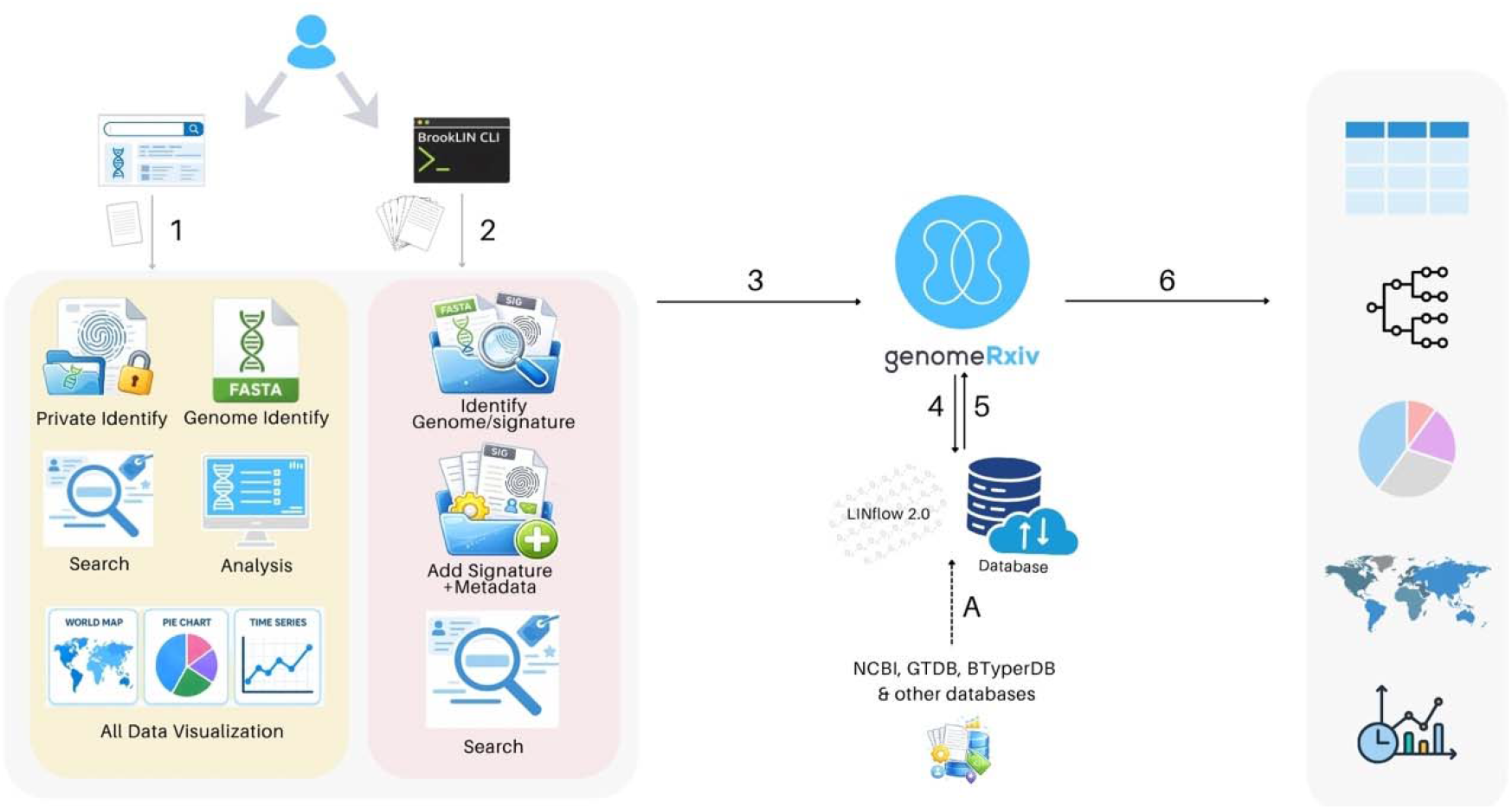
Overview of the *genomeRxiv* web server. The user can access the web server either through the the *genomeRxiv* website (1) or command line tool *BrookLIN* (2). Queries are passed to the *genomeRxiv* backend (3), which consists of a relational database into which data from NCBI, GTDB, and other databases are regularly imported (A) and in which every genome has a LIN assigned by the LINflow 2.0 pipeline. The database is accessed each time a user submits a genome or signature for identification (using the LINflow 2.0 pipeline) or through simple keyword or LIN searches (4,5). Finally, results are passed to the website to be visualized (8) as tables, trees, maps, and charts. Icons included in this figure were either generated using ChatGPT (GPT-5.3) or sourced from canva (Pro).

### The *genomeRxiv* web pages and the *BrookLIN* command line tool

The *genomeRxiv* web service presents the user with a number of pages, each with a defined role or function.

#### “Genome Identify” and “Private Identify” pages

The *Genome Identify* page serves as the landing (or *Home*) page for the user. No account is required but if users create an account (using the *Sign in* page accessible through the *Menu*) this enables them to track the history of their activities with links to saved results. The *Genome Identify* page accepts FASTA files or NCBI accession numbers as queries, and the *Private Identify* page accepts FASTA files or sourmash signatures as queries. On the *Genome Identify* page, clicking the *Identify* button triggers submission of the user’s FASTA file query to the *genomeRxiv* server (either by file upload or retrieval of the FASTA file corresponding to the entered accession number from NCBI). A temporary LIN is assigned to the query using the LINflow pipeline^32^, based on ANI to the most similar genome with a LIN in the *genomeRxiv* database. The temporary LIN is the basis for all results that will be displayed on the *Results* page (see below).

The *Private Identify* page assigns LINs without the need to share the genome sequence with the genomeRxiv service. Clicking the *Identify* button on this page instead triggers computation of a sourmash FracMinHash signature directly in the user’s browser. The generated signature is then uploaded to the *genomeRxiv* server and a LIN is assigned to it based on its ANI, inferred by sourmash, to the best hit in the *genomeRxiv* database. If the user submits a precomputed sourmash signature as the query, this signature is uploaded and used for LIN assignment. As with *Genome Identify*, the temporary LIN immediately provides identification of the query, placing it into the hierarchical taxonomic LIN framework. In this case however, the LIN is based only on ANI inferred from a FracMinHash signature comparison instead of ANIb making the right-most positions not directly comparable to other LINs in the *genomeRxiv* database that were assigned based on ANIb.

#### “Results” page

As soon as a query is completed, all results are displayed on the *Results page*. The *Results* page provides information about the query, the most similar genome in the *genomeRxiv* database, and about the LINgroups to which the query belongs **(Fig. 3)**. The user can click on any of the information displayed to the right of the LIN of the closest genome to open a new tab that will display metadata associated with that genome. The user can also click on any of the LINgroups to which the query belongs to learn more about these groups. Leveraging the strain typing aspect of LINs, the user can also select any prefix of the tentative LIN assigned to the query to learn more about the genomes that share the selected prefix with the query.

**Fig. 3.**
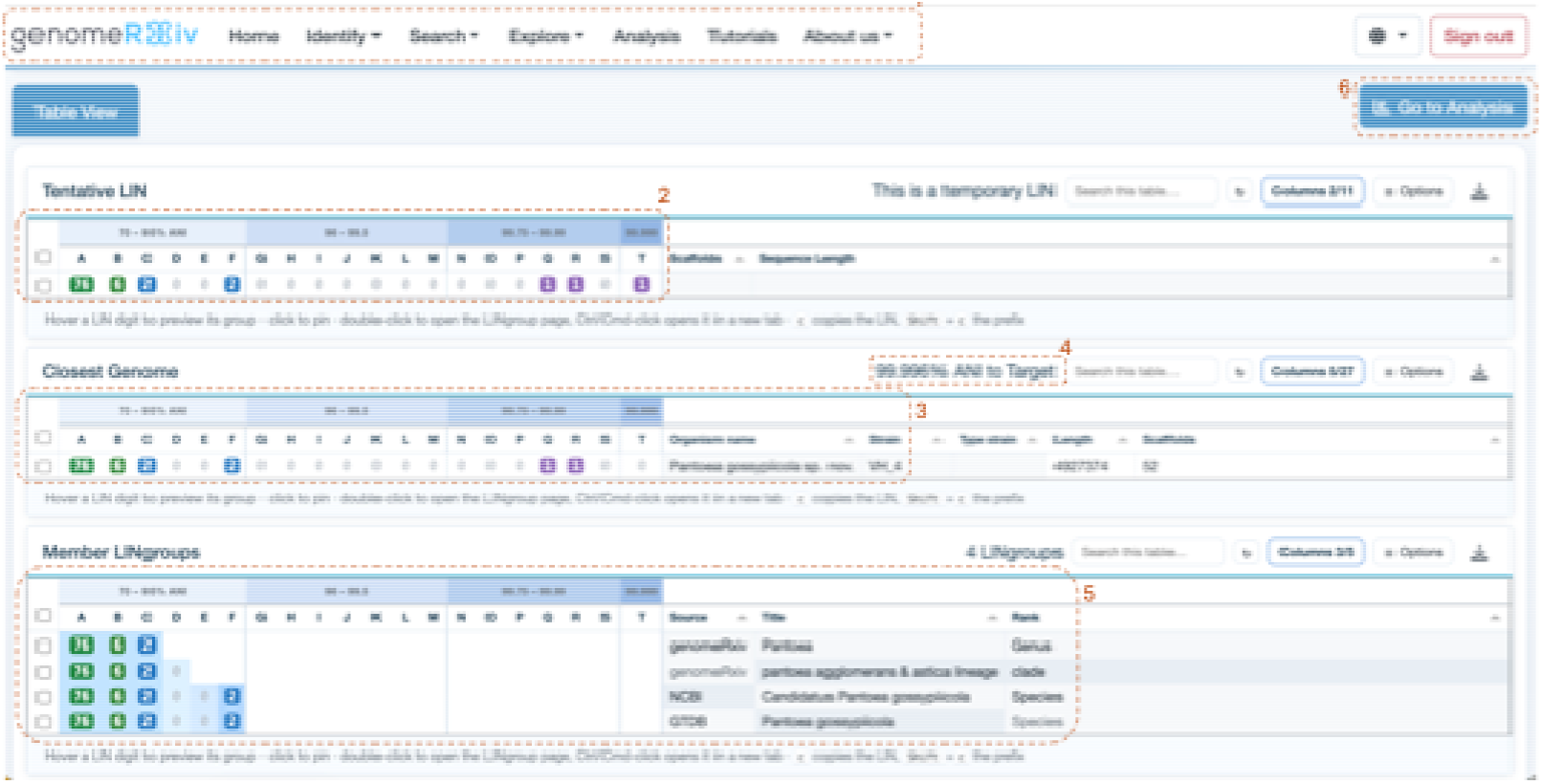
The *genomeRxiv* “*Result*” page. This page shows the result of “genome identify” and “private identify” queries. The key elements of the page are: **1** the navigation menu; **2** The tentative LIN assigned to the query providing strain typing information; **3** The LIN of the most similar genome. To the right is the organism and strain name associated with the closest genome and some additional information; **4** The ANI between the query and its closest genome in the *genomeRxiv* database; **5** the LINgroups to which the query belongs based on its tentative LIN providing taxonomic identification; **6** “Go to Analysis” button. Selecting one or more genomes and clicking this button opens the *Analysis* page, which allows the user to explore metadata of these genomes and to determine how they are related to each other based on a sourmash distance tree (see **Fig. 4**).

#### “Selected LINgroup” page

On the *Results* page, users can select any prefix shared between the query and the most similar genome by hovering over the displayed LINs. Once the user selects a prefix by double clicking on it or by clicking on any LINgroup name, the user is taken to the *Selected LINgroup* page. This page displays a list of described LINgroups that overlap with the selected prefix and presents buttons leading to three pages: *LINgroup genomes, LINgroup sketches*, and *Analysis*

#### “LINgroup genomes” page

This page lists all genomes that have a LIN which includes the user-selected prefix. These genomes are shown in a table in the *Table view* tab (**Fig. 4A**) by default; this table may be downloaded (optionally including all associated strain and genome metadata). The *Isolation Source* tab (**Fig. 4B**) summarizes the distribution of source categories for genomes based on associated NCBI metadata giving a hint at the ecology of the strains with the same LIN prefix. The *Collection Date* tab displays a timeline for collection of sequenced strains (where data are available in NCBI). The *Sampling Country* tab (**Fig. 4C**) shows the geographic distribution of sequenced strains by country (again, based on available NCBI metadata). The *Sampling Origin* tab provides more precise location data within countries, where available. The *Distance Tree* tab (**Fig. 4D**) renders a sourmash distance tree, which can be modified using a range of tree visualization options.

**Fig. 4.**
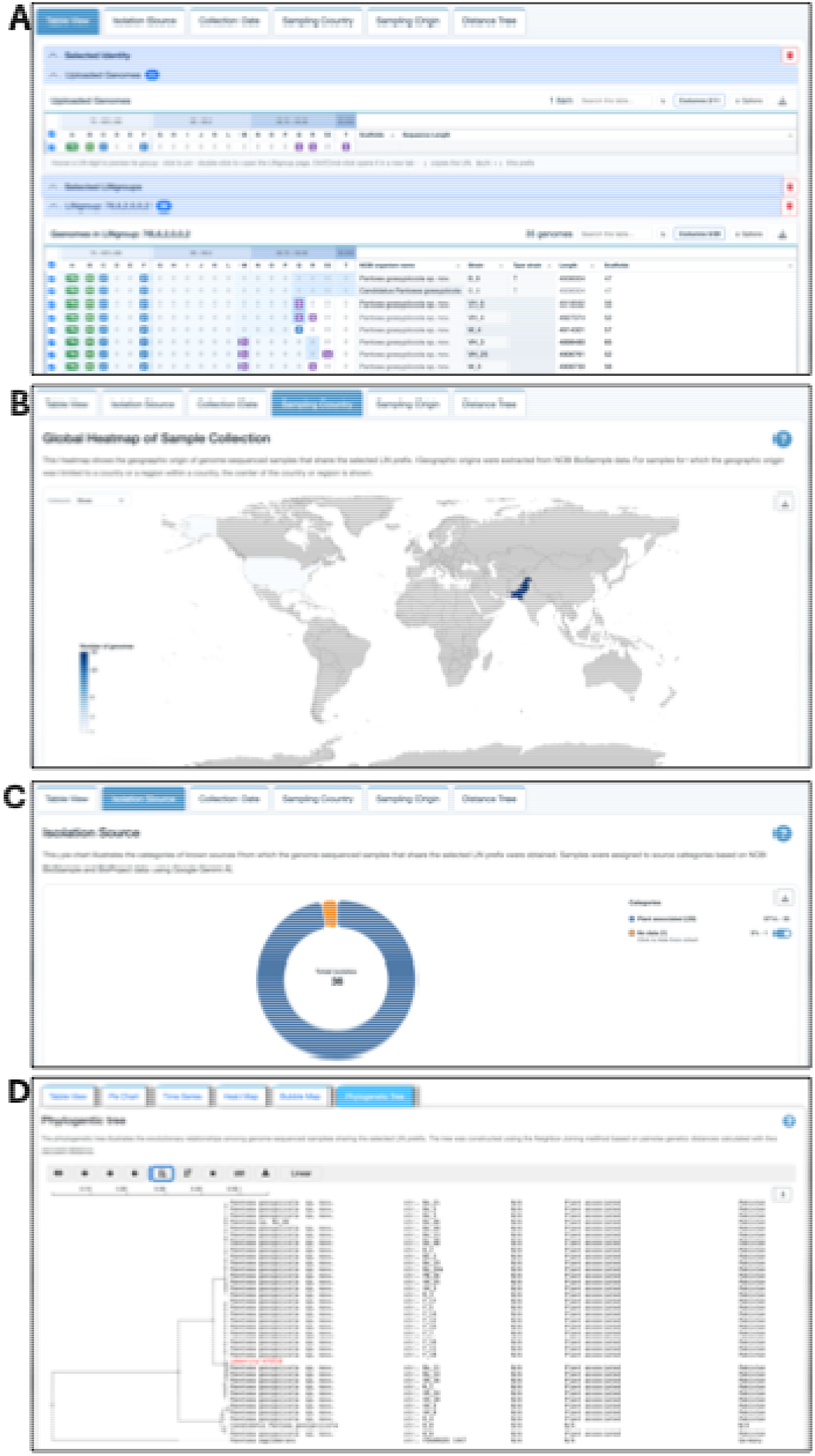
The *genomeRxiv* “*LINgroup genomes*” and “*Analysis*” pages. Both pages allow users to view genomes in table format (A), the geographic locations of where the genome-sequenced strains were isolated (B), the isolation source categories (C), and a sourmash distance tree (D).

#### “LINgroup sketches” page

This page is equivalent to the *LINgroup genomes* page but for user-submitted sourmash signatures and associated user-provided metadata (see *BrookLIN* section below). As few signatures have so far been uploaded to *genomeRxiv*, this page is sparsely populated for most LIN prefixes.

#### “Analysis” page

The *Analysis* page extends the functionality of the *LINgroup genomes* and *LINgroup sketches* pages by giving users the flexibility to include any number of query genomes and query signatures in combination with any selection of genomes and/or signatures from one or more LINgroups. The *Analysis* page was designed to address a user’s need to quickly visualize how their newly sequenced query genomes are related to each other and to already published genomes in NCBI without the need to build core genome trees, which requires substantial computing resources. When visualizing a tree of the selected genomes and/or signatures, users may find it useful to include a genome from a sister taxon in their selection so the tree can be rooted on an outgroup. Selecting a genome as root is done by simply clicking on the respective tip label in the tree and choosing “Reroot on this node” from the menu.

#### “Genome search” and “LINgroup search” pages

These pages allow users to browse *genomeRxiv* using keywords (species or strain names, isolation sources, and more) or LIN prefixes. The respective result pages are similar to the *Identify* result pages from which users can explore isolation sources, time series, geographic distribution, and distance trees for the selected genomes and/or LINgroups.

#### “Tutorials” and “About us” pages

Users can access short instructional videos on how to use *genomeRxiv* in the *Tutorials* menu and learn more about the LIN concept, funding sources, the *genomeRxiv* team, and other background information by clicking on the *About us* menu.

#### The BrookLIN command line tool

The *BrookLIN* command line tool supplements the website by allowing users to submit queries in batch mode and to allow developers of computational tools and pipelines to query *genomeRxiv* through its API. The respective commands trigger functions similar to the ones available on the website. One difference is that *Private Identify* is replaced by the *Signature Identify* command, which can only use signatures as queries. A key *BrookLIN* function, which is not available on the website, is that users can upload FracMinHash signatures and associated metadata to the *genomeRxiv* database to publicly share the precise identity of their sequenced genomes (since signatures are assigned public LINs and can be used for phylogenetic tree construction by other users) without sharing their actual genome sequences. Instructions on how to access and use BrookLIN can be found at https://code.vt.edu/linbaseproject/brooklin.

### Overview of genomeRxiv content

On October 1, 2026, the *genomeRxiv* database contained 354,448 bacterial and prokaryotic genomes. Users can view source categories, times of isolation, and geographic locations of isolations based on available NCBI metadata for all genomes included in the *genomeRxiv* database under the *Explore* menu at the *Isolation Source, Collection Date*, and *Geographic origin* links. The *genomeRxiv* database also includes a small number of sourmash signatures, and we anticipate the addition of many more signatures as users start using the *BrookLIN* command line tool to share signatures of yet unpublished genomes.

Regarding described LINgroups, *genomeRxiv* includes 153, 083 LINgroups corresponding to named species and numbered species clusters from GTDB release 10-RS226^33^ and NCBI (downloaded in March 2026. These LINgroups are circumscribed based on GTDB representatives and NCBI genomes labeled as type strains, respectively, using the ANI radius specified by GTDB for GTDB species and a default 95% ANI threshold for NCBI species. In cases where two GTDB species or two NCBI species were assigned the same prefix following this process, we extended the species prefix to the first position at which they diverged. We also included *Bacillus cereus* group circumscriptions from BTyperDB^34^, including *Bacillus cereus* subspecies *anthracis*. Finally, *genomeRxiv* includes LINgroups corresponding to named infraspecies lineages that we circumscribe based on published papers.

We invite experts on groups of organisms to share additional infraspecies descriptions of additional named infraspecies LINgroups. Upon request, users can get access to do this for individual circumscriptions directly on the *genomeRxiv* website by simply selecting the prefix to which they want to add a circumscription. We also provide an Excel template (**Supplementary Table 1**) for users to provide batches of circumscriptions by email.

### Examples of identifying a genome or signature

As an example of how *genomeRxiv* can identify an important human pathogen at infraspecies resolution, we use the public genome assembly (GCF_000590755.1) of strain J/31-98 of Chlamydia trachomatis, the causal agent of Chlamydia and trachoma diseases.

Fig. 5 shows the Genome Identify result at species (GTDB and NCBI) and lineage resolution and a tanglegram reveals how the distance tree generated in seconds on the genomeRxiv website aligns with an 873-gene core genome tree of the same genomes at the lineage level (ANI of 99.25-99.75) and how most individual genomes belong to similar clades in both trees. The main differences between the trees are in clade order but not in clade content.

**Fig. 5.**
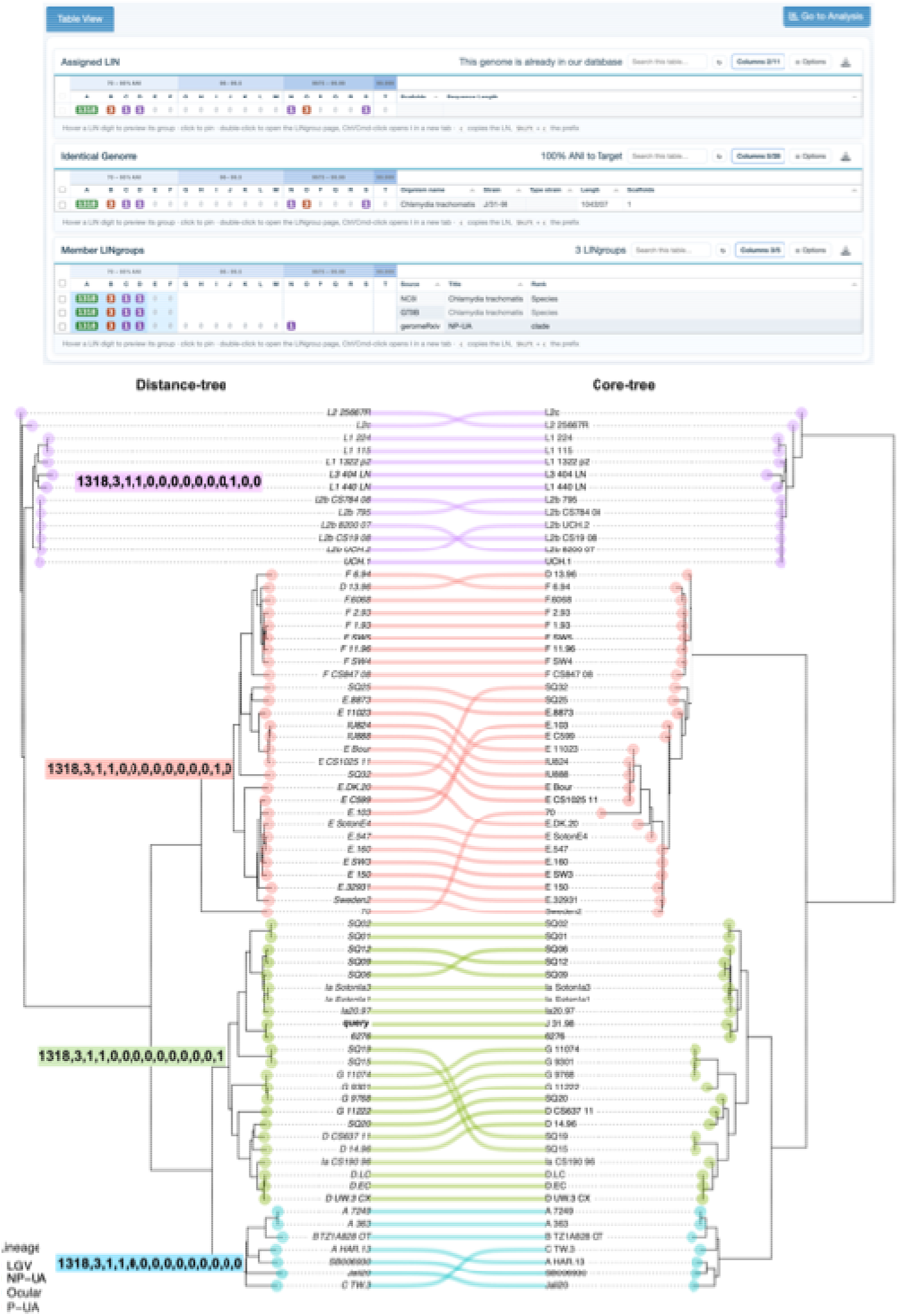
*Genome Identify* result. **A** *Genome Identify* results for *Chlamydia trachomatis* J/31-98 is shown at the top. Since the genome was already in the genomeRxiv database, it has a 100% match to itself. **B** Comparison of the sourmash distance tree generated by the *genomeRxiv* website with a core genome tree of the query, Chlamydia trachomatis J/31-98, and related strains.

As an example of the *Private Identify* function, we use a genome of the biotechnologically important genus *Streptomyces*. The strain *Streptomyces* sp. PSAA01 (GCF_021890875.1) was reported in 2022 to produce the new antibacterial compound pyrimidomycin^35^. Assuming the authors had been required to keep the identity of this strain private, they could have used the *Private identify* function to compute a sourmash signature in their browser. This would have allowed them to identify their genome as a member of the species *S. yatensis* and to determine how this strain is related to the type strain of this species and to strains of related species (**Fig. 6**,). As with C. trachomatis, the distance tree generated in near real-time on the “*Analysis*” page (left) is identical to a 1979-gene core genome tree (right) with the only difference being a single switch in clade order.

**Fig. 6.**
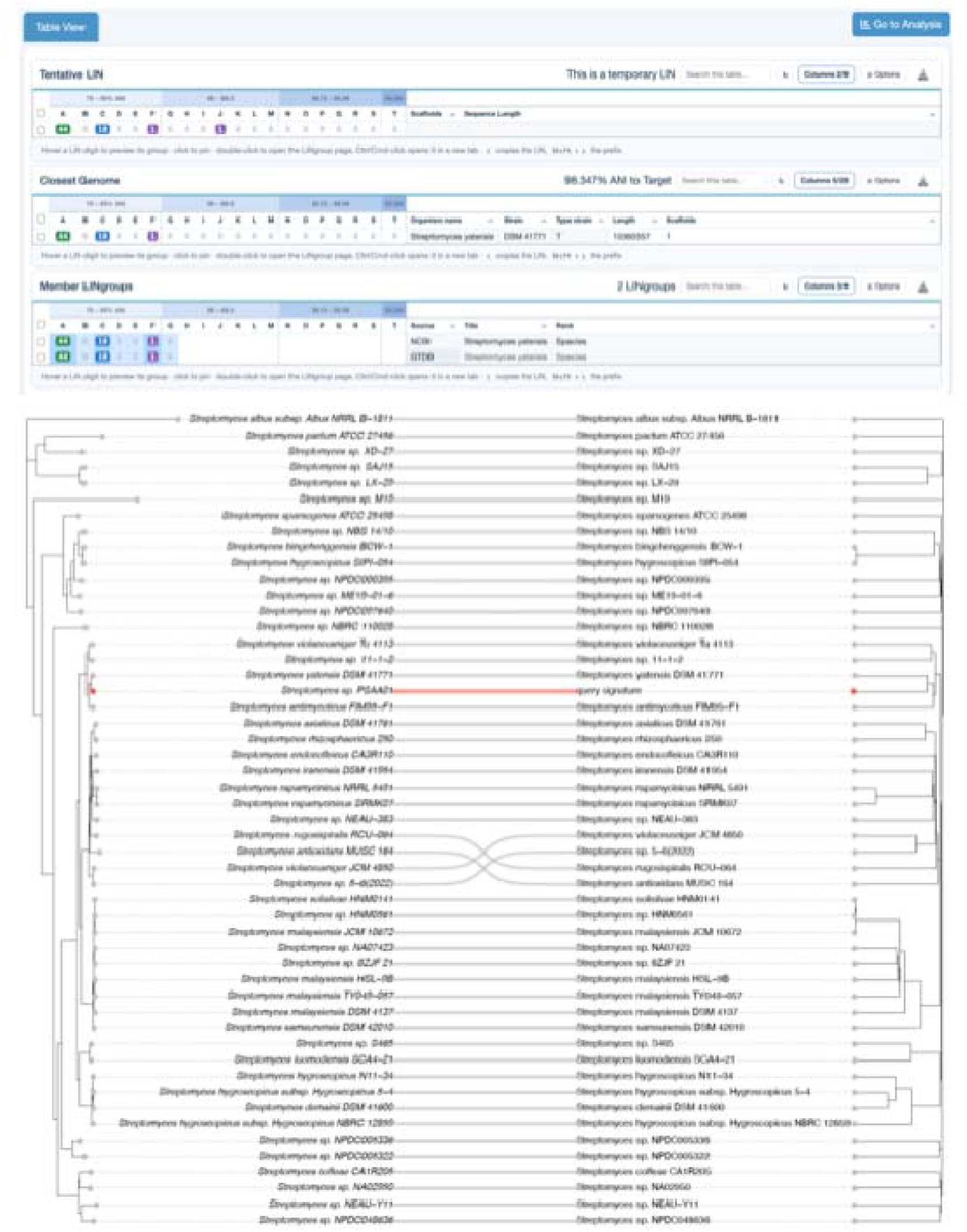
Private Identify result and comparison of the sourmash distance tree generated by the *genomeRxiv* website with a core genome tree of the query, *Streptomyces* sp. PSAA01, and related strains.

## Discussion

We describe *genomeRxiv*, a web server that bridges the gap between prokaryotic taxonomy and strain typing permitting genome-based identification of almost any prokaryotic genome at near-strain level resolution. This is accomplished by expanding use of ANI from the single species rank threshold to a hierarchical framework of 20 ranks of ANI thresholds starting approximately at the genus rank and approaching the single strain level.

*genomeRxiv* can be used for precise identification of isolates of any emerging and/or novel strain of any bacterial pathogen in the realm of disease surveillance and biosecurity without the need to set up a pathogen-specific typing scheme. This distinguishes *genomeRxiv* from pathogen-specific typing infrastructures such as GenomeTrakr^10^, Enterobase^11^, BIGSdb Pasteur^12^, or PhytoBacExplorer^36^, which set up an *ad hoc* scheme for each pathogen and which do not cover the majority of human, animal, and plant pathogens, nor any non-pathogenic prokaryote. *genomeRxiv* is also conceptually different from other existing web services for taxonomic identification, such as TYGS^9^ and GTDB-toolkit^8^, which are limited to the species as the smallest unit.

By assigning a unique LIN to every individual genome, *genomeRxiv* accomplishes two goals. First, the assigned LIN is informative of how similar that genome is to all other related genomes, from which one can immediately infer evolutionary relationships. When new genomes are added, the LINs of all genomes that already have a LIN do not change and still reflect evolutionary relationships to the newly added genomes. Second, the LIN assigned to a newly sequenced genome, and the LIN prefix this genome shares with the most similar genomes, provide identifiers for the newly sequenced genome and for the group of closely related genomes it belongs to, respectively. LINs thus provide a stable identifier scheme prior to, and independent of, official nomenclature or taxonomic opinion concerning any new lineage or new species. This persistent scheme facilitates communication about any new outbreak strain or emerging pathogen species.

A second key functionality of *genomeRxiv* is that LINs can be used to circumscribe any group of genomes between the genus rank and the strain level that are monophyletic, such as species complexes, named species, infraspecific phylotypes, or recently emerged genetic lineages. Multiple schemes can be recorded and reported simultaneously in genomeRxiv, even if they are in contradiction. For example, we have circumscribed both “NCBI species” and “GTDB species” within genomeRxiv, meaning a user can immediately see any discrepancies between these two important taxonomic resources. Several such classifications have been included in *genomeRxiv*, as exemplified by species and subspecies circumscriptions derived from BTypeDB^34^ for the *B. cereus* group, which were incorporated to address the need to precisely circumscribe high-impact, anthrax-causing pathogens from closely related, harmless *B. cereus* strains.

Since ANI becomes uninformative of evolutionary relationships above the genus rank, *genomeRxiv* cannot classify or assign LINs to bacteria at levels above the genus rank. A taxonomic tool such as the GTDB toolkit is more appropriate for these cases^8^. At the other extreme, *genomeRxiv* results may deviate from core genome phylogenies and cgMLST-based LIN codes at the right-most LIN positions. This can be due to the following factors. First, core genome trees and cgMLST-based LIN codes are by definition based on the alignment of orthologous genes shared by all genomes of a species (or even a group of related species). A LIN is instead assigned to a genome by computing the ANIb between this genome and its most similar genome in the database at that time. The alignable fraction between the two genomes that is used to compute ANI may be significantly shorter or longer than the core genome of a species and may include regions of the flexible genome that are less conserved than genes of the core genome. The alignable fraction may even include non-vertically inherited insertion sequences, transposons, and plasmids with pairwise identity as low as 30%, which is the threshold used by ANIb as implemented in pyani^37^, the program used in the LINflow LIN assignment pipeline^26^. Therefore, if *genomeRxiv* identifies genomes that appear to belong to the same outbreak and it becomes necessary to investigate that outbreak, it may still be necessary to develop a pathogen-specific cgMLST-based scheme and/or use SNP-trees to infer transmission chains. For the same reasons, *genomeRxiv* is intended to complement, not to replace, validated cg-MLST schemes and LIN codes^15^ that already exist within the Enterobase^11^ and BIGSdb Pasteur^12^ platforms.

Another key strength of *genomeRxiv* compared to other web servers is the ability to use *k*-mer signatures (sourmash FracMinHashes) instead of genome sequences. Signatures can be used as queries (either computed with sourmash by the user or computed in the browser by *genomeRxiv*) and be uploaded to the databases through the *BrookLIN* command line tool. Other users can then find these signatures and compare them to their own signatures without the need to share or use the genome sequence itself. This allows users to keep their actual genome sequences private while still sharing their precise identity. This may be important for users to comply with the Nagoya protocol^30,31^, so that the benefits from genes in a bacterial genome, for example those encoding novel antibiotics, do not become accessible before guaranteeing that the benefits are shared with the country of origin. Similarly, a company that sequenced thousands of bacterial genomes may be looking for academic collaborators without wanting to release the genome sequences themselves because of concerns about intellectual property protection. Finally, signatures uploaded by users, in combination with the signatures of public genome sequences already included in *genomeRxiv*, could be used to detect human, animal, and plant disease outbreaks. In fact, it was recently shown that sourmash FracMinHash sketches have sufficient resolution as a first step in genomic epidemiology^38^. *genomeRxiv* visualisations of source categories, times of isolation, geographic locations, and signature-based distance trees can further support epidemiological investigations and give insights into ecological niches and geographic prevalence of environmental bacteria and archaea. This has important regulatory applications, since these visualizations (in combination with the downloadable metadata), can support applications for field releases or commercialization of biologicals, for which USDA APHIS PPQ requires biotech companies to provide prevalence data^39^.

Several limitations of the current *genomeRxiv* version still require to be addressed. Not all public genomes from NCBI have yet been included and we have not yet set up automated updates to include new genomes as they are released by NCBI. Also, updates to species circumscriptions based on future GTDB releases still need to be initiated manually. We are planning to completely automate such updates, pending future funding. Named infraspecies groups have only been circumscribed in *genomeRxiv* for a small number of species, and we depend on contributions by species-specific subject experts to provide additional circumscriptions, although AI agents are likely to be able to perform automated literature searches and take over some of the manual labor in the near future. Similarly, because many genomes in NCBI are missing key metadata about isolation sources, adding such metadata based on literature searches would greatly improve the information a user can gain from *genomeRxiv*. Also, due to hardware restrictions, *genomeRxiv* currently has limited storage capacity and computing power, limiting speed and number of concurrent users it can handle compared to NCBI Genbank or the JGI genome portal. We envision that in the long term *genomeRxiv* may continue as an improved, more powerful, stand-alone web server, possibly funded by foundations, similar to the bioRxiv preprint server, or through user fees charged to commercial and government entities. Alternatively, *genomeRxiv* capabilities could be incorporated into NCBI Genbank and/or the JGI Genome Portal, or other government-funded services.

In conclusion, *genomeRxiv* shows how a strictly genome-similarity based classification approach in combination with FracMinHashes can bridge the gap between taxonomy and strain typing and offer a universal solution for near real-time and near strain-level identification of any prokaryote, with applications ranging from human, animal, and plant epidemiology to biosecurity and biotechnology.

## Methods

### LIN assignment and queries

LINs are assigned to genomes offline using the LINflow 2.0 pipeline^32^. Importantly, LIN assignment is not recomputed for every query; newly added genomes are assigned incrementally and appended to the existing hierarchy. Genome similarity search is accelerated with sourmash^27,40^ FracMinHash sketches (*k*-mer sizes 21 and 51; scaled factor 1000), indexed for rapid candidate retrieval. This staged design reflects a practical tradeoff from large-scale genome analysis: alignment-based comparisons are biologically interpretable but expensive, whereas sketch-based comparisons are fast but approximate. *genomeRxiv* uses sketches for high-throughput candidate retrieval and applies ANIb refinement where sequence-level resolution is required. For sequence-based queries, candidate hits are refined with BLAST-based ANI (ANIb) using *pyani^28^* before final LIN assignment.

The three user-facing query modes differ mainly in where sketches are computed and what is uploaded: *Genome Identify*: FASTA or accession input’ sequence-based workflow on server, temporary LIN plus LINgroup reporting; *Private Identify*: FracMinHash sketch computed client-side in browser, only sketch is uploaded (not the query genome sequence); *BrookLIN Signature Identify*: precomputed signatures are submitted through CLI for batch and automated workflows. On average, assigning a query genome to a LIN takes approximately 1–2 minutes on the website, with most runtime spent on ANIb refinement of top candidates. Runtime increases as the number of genomes available in clusters at the 70% ANI threshold grows.

### The *genomeRxiv* database and its content

The *genomeRxiv* database stores genome identity, LIN assignments, signatures, LINgroups, and metadata in a relational schema (MySQL). Core entities include: genome records (accession, organism name, source metadata, quality fields), LIN assignments (scheme identifier, full LIN, prefix-indexed fields), sketch/signature records (FracMinHash signatures and provenance), LINgroup definitions (name, taxonomy source, circumscription method), and LINgroup-prefix mapping tables used for fast membership lookup. Metadata are integrated from NCBI and GTDB snapshots, normalized to a common internal representation, and linked to both genome and LINgroup records. One sentence on how we categorize source categories with a link to a GitHub page with code repository. Prefix and LINgroup queries are index-backed so that query interpretation can be returned quickly during interactive use. See **Supplementary Figure 1** for an entity-relationship diagram of the *genomeRxiv* database schema, illustrating the core tables and their relationships.

### LINgroup circumscriptions and membership

LINgroups are circumscribed in two complementary ways. With ‘representative-radius mode’, a representative genome and an ANI radius are selected. The LIN prefix of the LINgroup is then determined by applying the radius to the LIN of the representative genome and selecting the corresponding length of the prefix. With ‘prefix mode’, LINgroups are defined directly by one or more prefixes (**Supplementary Table 1**). For a query with temporary LIN or when looking up member genomes or signatures of a LINgroup, LINgroup membership is determined based on prefix matching.

### The *genomeRxiv* web site

The *genomeRxiv* website exposes identification, browsing, and analysis workflows through a single web interface connected to the LIN, LINgroup, and signature backend. Interactive pages share common query services so that results are consistent across *Genome Identify, Private Identify, Search*, and *Analysis* views. Visualization modules are integrated with database-backed filtering so users can move from a query result to subgroup summaries and distance trees without exporting intermediate files. The Javascript package phylotreeJS was used for tree construction^41^.

### BrookLIN command line tool

The BrookLIN command line tool mirrors core website functions for scripted and batch execution. Compared to browser workflows, BrookLIN is intended for reproducible pipeline integration and high-throughput use in surveillance and operational settings. For most BrookLIN commands, the workflow is input submission, server-side analysis, and result retrieval. The command-specific mappings are: *genome-identify*: sends a genome assembly for server-side identification and returns a tentative LIN, the closest reference genome, and matching LINgroups; *signature-identify*: sends a precomputed sourmash signature for privacy-preserving identification and returns tentative LIN placement plus matching genomes/LINgroups; *add-sketch*: uploads a signature and optional metadata so it can be stored and made discoverable in genomeRxiv; *genome-search*: queries stored genome records using LIN, taxonomy, and metadata criteria, then returns matching genomes; *lingroup-search*: queries described LINgroups using LIN prefixes, keywords, and describer fields, then returns matches; *batch*: applies one selected BrookLIN mode to many inputs, reusing the same backend services as the chosen mode.

### Comparison of LIN-based trees and core-genome trees

For each of the examples illustrated, the LIN-trees were downloaded from genomeRxiv as a newick tree file. The corresponding core-tree was constructed after performing core-genome analysis using PIRATE (version 1.0.5)^42^, following gene annotations performed with BAKTA (version 1.9.2)^43^ under default settings. The resulting core gene alignment was used as input for *IQ-TREE2* (version 2.3.0)^44^ to infer a maximum-likelihood phylogenetic tree using automated model selection. The final tanglegrams were constructed using the *phytools* package^45^ in R.

## Data availability

Source code, deployment assets, and CLI resources are publicly available at the following URLs: genomeRxiv web platform: http://genomerxiv.cs.vt.edu/, BrookLIN CLI: https://code.vt.edu/linbaseproject/brooklin, LINflow pipeline used by genomeRxiv: https://code.vt.edu/linbaseproject/LINflow

## Acknowledgements

We thank the following Virginia Tech Computer science undergraduate students who helped with various aspects of genomeRxiv during its development even if their code was not included in the final product: Abhilash Chauhan, Aman Kothari, Atul Bharadwaj, Chandra Sekhar Nerella, Matthew Russell, Pradyumann Singhal, Rahul Ramarao, and Suvasish Pant.

## List of Supplementary Files

Supplementary Table 1. Template to circumscribe infraspecies LINgroups.

**Supplementary Figure 1**. *genomeRxiv* database schema.

