## Supplementary figures and images for "genomeRxiv: a genome-based identification web server that bridges the gap between taxonomy and strain typing"

### Supplementary Figure 1

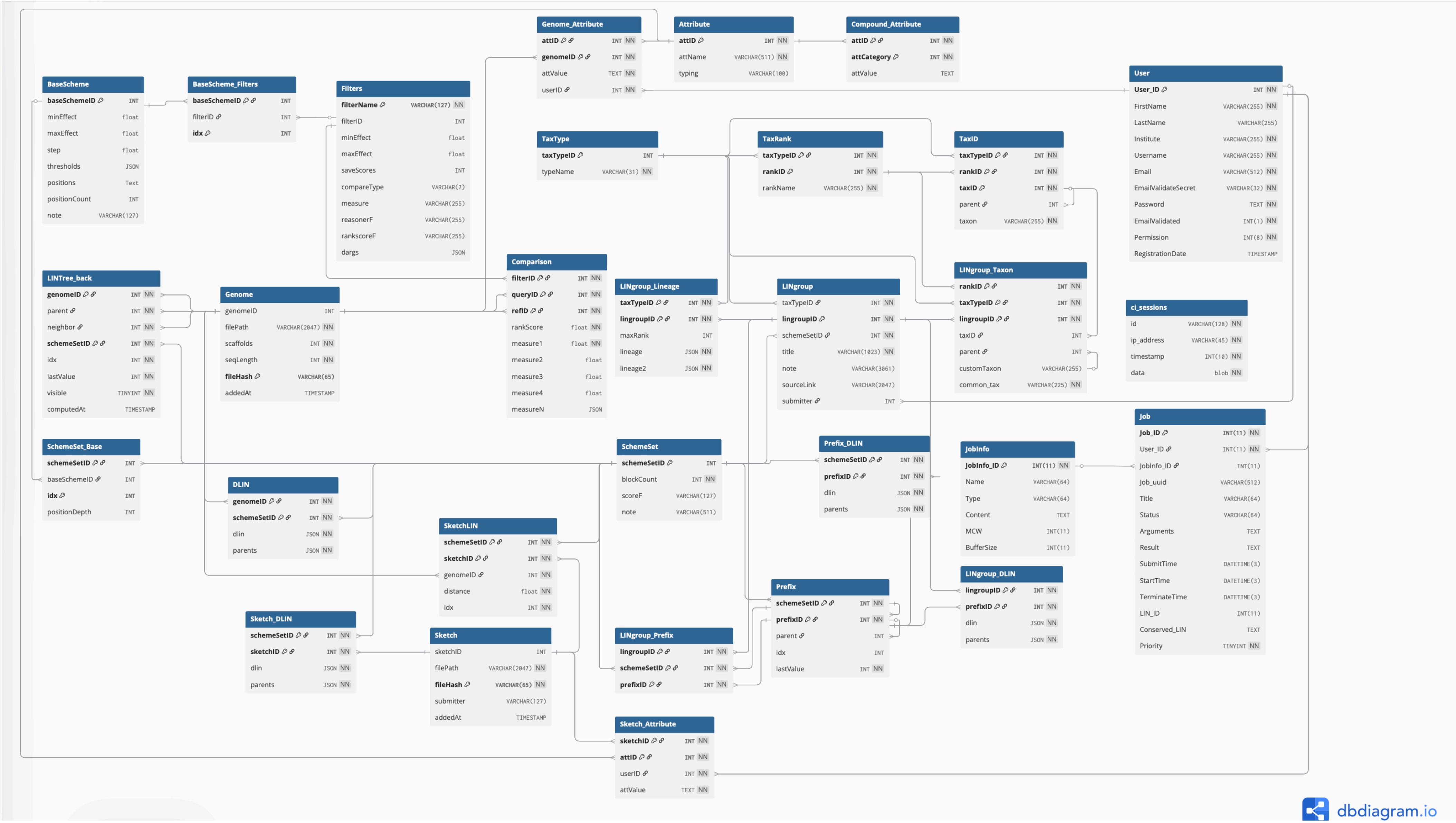
